# WNT signaling in human pluripotent stem cells promotes HDAC2-dependent epigenetic programs and development of retinoic acid-responsive mesoderm

**DOI:** 10.1101/2025.06.06.657928

**Authors:** Bao Q. Thai, Stephanie A. Luff, Jared M. Churko, Jonathan N. Young, Christopher M. Sturgeon, Deepta Bhattacharya

## Abstract

Human pluripotent stem cells (hPSCs) can be used as a scalable source of lymphocytes for adoptive cell therapies, contingent on the robust generation of definitive hematopoietic intermediates. Early WNT activation with CHIR99021 during mesoderm induction promoted the formation of KDR+ ALDH1A2+ mesodermal progenitors and subsequent generation of T cells in a retinoic acid (RA)-dependent manner. Integrated scRNA-seq and ATAC-seq defined a WNT-dependent developmental trajectory from hPSCs to KDR+ ALDH1A2+ mesoderm. Gene regulatory network modeling predicted HDAC2 and E-box transcription factors as regulators of RA-responsive mesodermal differentiation downstream of WNT. HDAC2 knockout impaired, while HDAC2 overexpression enhanced, KDR+ ALDH1A2+ progenitor formation. E-box factor manipulation had no discernible effect. An orthogonal chemical screen confirmed that HDAC2 inhibition suppressed KDR+ ALDH1A2+ mesodermal progenitors, whereas modulating histone methylation enhanced their formation. These findings reveal mechanisms by which WNT signaling promotes RA-responsive mesoderm and suggest methods to improve the generation of lymphocytes from hPSCs.

## Introduction

Human pluripotent stem cells (hPSCs), including human embryonic stem cells (hESCs) and induced pluripotent stem cells (iPSCs), offer a scalable starting point for allogeneic adoptive lymphocyte therapies to treat cancer and autoimmune diseases (Cichocki et al., 2020; Nishimura et al., 2013; Themeli et al., 2013). hPSCs have unlimited self-renewal potential and, with the advent of targeted nuclease technology, are amenable to genetic engineering (Hockemeyer and Jaenisch, 2016). Genetic modifications in hPSCs, such as chimeric antigen receptor (CAR) integration for tumor targeting, can be stably maintained through differentiation (Themeli et al., 2013). Moreover, edits to ablate major immune recognition pathways may enable hPSC-derived cells to evade rejection and serve as a source for off-the-shelf allogeneic therapies (Gornalusse et al., 2017; Pizzato et al., 2024; Xu et al., 2019).

A critical bottleneck, however, remains the inefficient generation of hematopoietic progenitors with full lymphoid potential from hPSCs. Lessons from both primary human fetal hematopoiesis and hPSC modeling have demonstrated that multiple progenitors with lymphoid potential emerge during development, but many of these intermediates do not yield lymphocytes robustly (Luff et al., 2024). An intermediate population of KDR+ ALDH1A2+ mesoderm is associated with the emergence of retinoic acid (RA)-dependent definitive hematopoiesis and robust lymphopoiesis (Luff et al., 2022). These RA-responsive progenitors are critically dependent on WNT signaling during mesoderm differentiation (Luff et al., 2022; Sturgeon et al., 2014).

In this study, we examined the signaling, transcriptional, and epigenetic pathways downstream of WNT signaling that promote RA-responsive mesodermal progenitor differentiation from hPSCs. Using CHIR99021, we confirm that early WNT activation in embryoid bodies generates KDR+ ALDH1A2+ mesodermal progenitors that give rise to T cell progenitors in an RA-dependent manner. Multiome single-cell RNA- and ATAC- sequencing, pseudotime trajectory analysis, gene regulatory network modeling, and genetic manipulations identified HDAC2 as a regulator of RA-responsive mesoderm emergence downstream of WNT. A targeted small-molecule screen of epigenetic inhibitors revealed that selective modulation of histone methylation enhances progenitor output. These findings help provide an epigenetic and transcriptional explanation for how to generate RA-responsive mesoderm, an intermediate progenitor essential for subsequent definitive hematopoiesis.

## Results

### WNT signaling induces the generation of RA-responsive mesodermal progenitors capable of definitive hematopoiesis in hPSCs

In our previous studies, we identified that the emergence of a small but distinct ALDH1A2*+* population within early mesoderm correlated with the emergence of RA-dependent definitive hematopoiesis, and that this population was specified in a WNT-dependent manner (Luff et al., 2022). Manipulating BMP4 and FGF concentrations did not markedly impact the numbers of these cells (data not shown). Recent protocols to generate multipotent hematopoietic stem and progenitor cells from hPSCs activate WNT signaling from the very beginning of mesoderm induction (Fowler et al., 2024; Ng et al., 2024). Therefore, to improve the efficiency of generating this population from hPSCs, we tested the impact of early WNT signal activation on the yields of ALDH1A2+ mesoderm. We treated hPSCs with CHIR99021, a glycogen synthase kinase 3 beta (GSK3β) inhibitor that activates canonical WNT signaling via stabilization of β-catenin, from day 0 (early CHIR) or day 2 (late CHIR) during a three-day embryoid body (EB) differentiation protocol. On day 3, we assessed the emergence of RA-responsive mesodermal progenitors by flow cytometry using KDR as a mesodermal marker and ALDEFLUOR as a reporter of the RA-processing enzyme aldehyde dehydrogenase (ALDH) (gating strategy in **Figure S1A**). Flow cytometric quantification showed that CHIR99021 significantly increased the generation of KDR+ ALDEFLUOR+ mesodermal progenitors on day 3 (**Figures 1A-B**), and CD34+ hematopoietic progenitors on day 16 of EB cultures (**Figures S1B-C**), derived from both H1 hESCs and an iPSC line derived from a human tonsillar B cell. To confirm that these KDR+ ALDEFLUOR+ and subsequent CD34+ progenitors were correctly specified for definitive hematopoiesis, we performed T cell assays (gating strategy in **Figure S1D**). When RA was added at Day 3, early CHIR cultures robustly yielded CD7+ NK/T progenitors and T-committed CD5+ CD7+ progenitors, while late CHIR cultures did not (**Figure 1C**). Excluding RA from early CHIR cultures markedly reduced T-committed progenitors from both H1 hESCs and iPSCs (**Figure 1D**). These results indicate that early WNT signaling promotes the development of RA-responsive mesoderm progenitors important for the subsequent formation of definitive hematopoietic progenitors.

**Figure 1.**
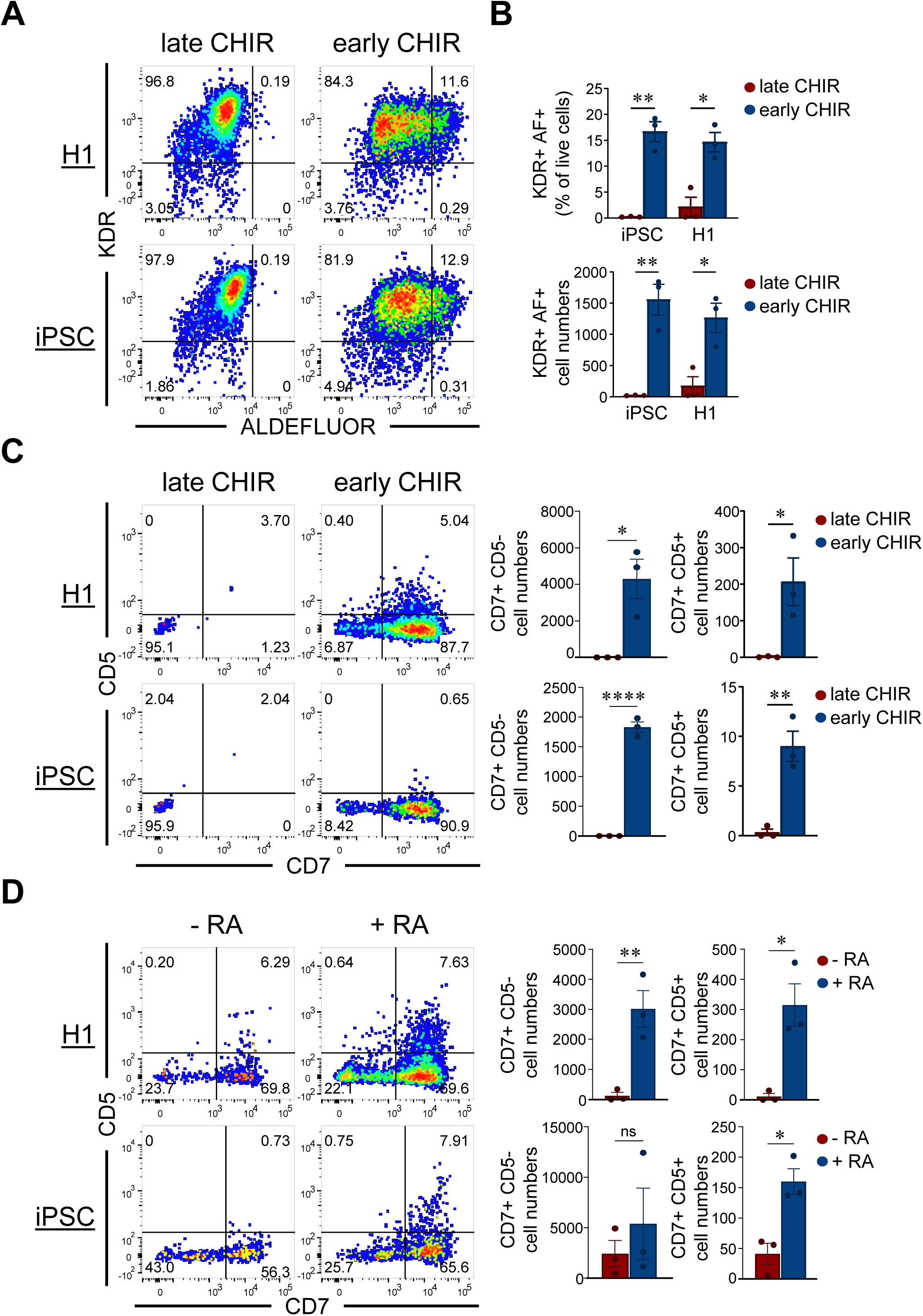
WNT signaling induces the generation of RA-responsive mesodermal progenitors capable of definitive hematopoiesis in hPSCs. (A) Representative flow cytometry plots showing expression of KDR and ALDEFLUOR in H1- and iPSC- derived mesodermal progenitors on day 3 of EB differentiation, with CHIR99021 added on day 0 (early CHIR) or on day 2 (late CHIR). (B) Quantification of the percentage of KDR+ ALDEFLUOR+ cells (top) and total KDR+ ALDEFLUOR+ cell numbers (bottom) in H1- and iPSC-derived cultures. Statistical analysis was performed using 2-way ANOVA with Šídák multiple comparisons test and *p < 0.05, **p < 0.01. Data are presented as mean ± SEM. Each circle represents an independent experiment. (C) Representative flow cytometry plots showing the expression of CD5 and CD7 in H1- and iPSC-derived T cell progenitors on day 14 of T cell differentiation from +/- CHIR progenitors treated with RA on Day 3 (left panels). Quantification of the total number of CD7+ CD5- T/NK progenitors and CD7+ CD5+ T-committed progenitors in H1- (top right) and iPSC-derived cultures (bottom right). Statistical analysis was performed using 2-tailed t-test, with *p < 0.05, **p < 0.01, ****p < 0.0001. Data are presented as mean ± SEM. Each circle represents an independent experiment. (D) Representative flow cytometry plots showing the expression of CD5 and CD7 in H1- and iPSC-derived T cell progenitors on day 14 of T cell differentiation from untreated and RA-treated progenitors (left panels). Quantification of the total number of CD7+ CD5- T/NK progenitors and CD7+ CD5+ T-committed progenitors in H1- (top) and iPSC-derived culture (bottom). Statistical analysis was performed using two-tailed t-test, with *p < 0.05, **p < 0.01. Data are presented as mean ± SEM. Each circle represents an independent experiment.

### WNT signaling induces transcriptional changes associated with definitive hematopoiesis

To define how early WNT signaling promotes KDR+ ALDH1A2+ mesodermal progenitor formation, we performed combined single-cell RNA sequencing (scRNA-seq) and single-cell ATAC sequencing (scATAC-seq) analysis on H1 hESCs during each of the first three days of differentiation. Analysis of integrated scRNA- and scATAC-seq data identified 18 distinct clusters (**Figure 2A**). Compared to late CHIR cells, early CHIR cells diverged transcriptionally and epigenetically by Day 1.5 (**Figure 2B**). ALDH1A2 expression was seen primarily in KDR+ PDGFRA+ mesoderm progenitors in early CHIR cultures on Day 3, corroborating our flow cytometric data (**Figure 2C-D**). Expression of ALDH1A1 and ALDH1A3 was minimal (**Figure S2A**). HOXA gene expression (HOXA9 and HOXA10) was observed in ALDH1A2+ cells on day 3 of early CHIR cultures, further supporting the role of WNT signaling in early formation of definitive hematopoietic progenitors (**Figure S2B)** (Dou et al., 2016; Ng et al., 2016).

**Figure 2.**
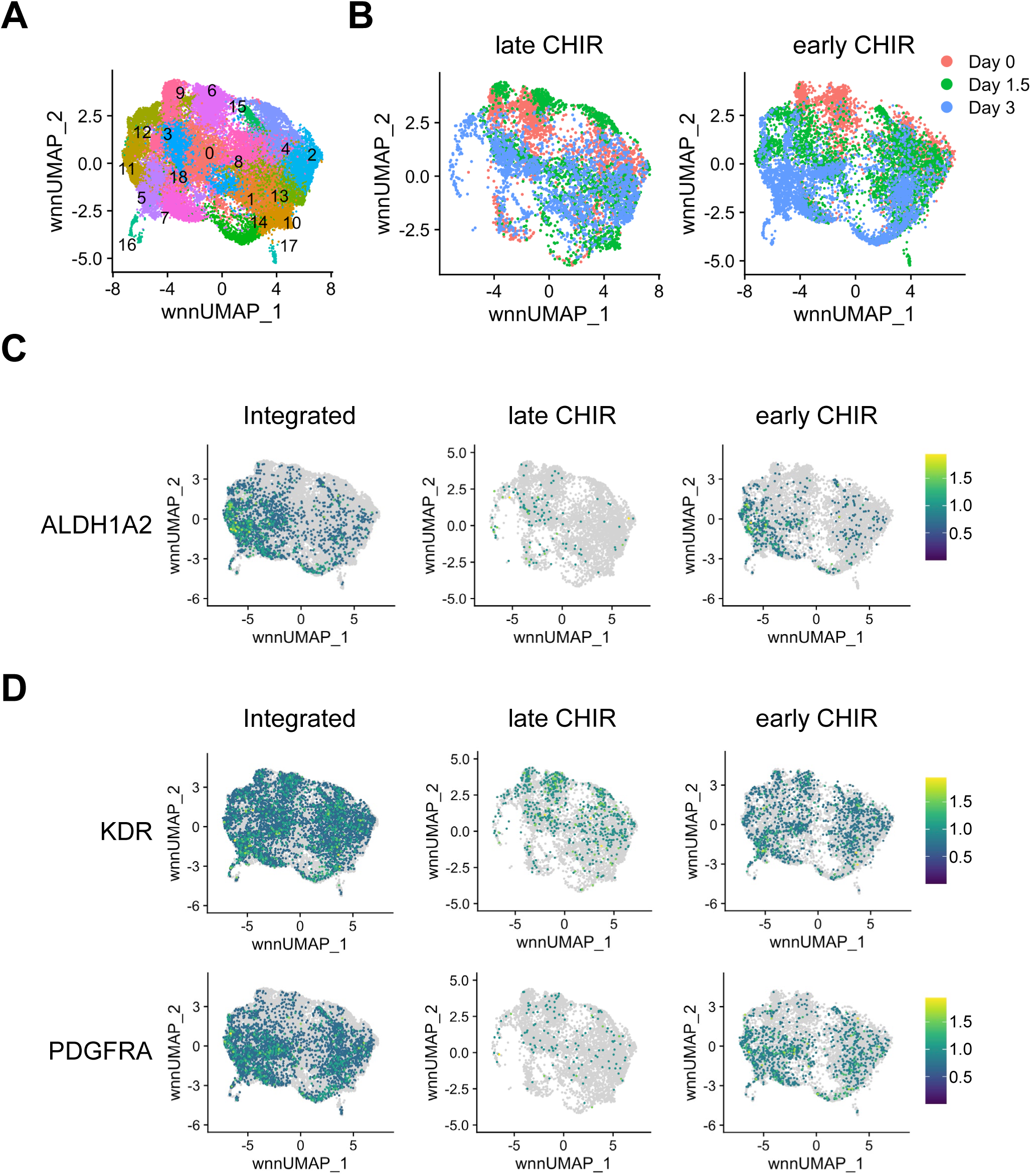
WNT signaling induces transcriptional changes associated with RA-responsive mesoderm formation. (A) UMAP plot of combined single-cell ATAC- and RNA-seq data integrated using the Weight Nearest Neighbor (WNN) method, showing distinct clusters of cells during the first 3 days of EB differentiation with CHIR99021 added on day 0 (early CHIR) or day 2 (late CHIR). (B) UMAP plots showing the progression of differentiation over time for early CHIR and late CHIR conditions. (C) Relative expression of ALDH1A2 in an integrated UMAP or separated as early CHIR or late CHIR conditions. Color bar is based on scaled relative expression. Gray indicates no detectable expression. (D) Relative expression of mesoderm markers *KDR* and *PDGFRA* in an integrated UMAP or separated as early CHIR or late CHIR conditions.

### In silico gene perturbations identified transcriptional and epigenetic regulators in the development of ALDH1A2+ KDR+ RA-responsive mesodermal progenitors

To identify regulators of the development of WNT-induced ALDH1A2+ *KDR*+ mesodermal progenitors, we reconstructed our UMAPs to focus only on early CHIR cells (**Figure 3A and 3B**). Using Monocle 3.0 (Cao et al., 2019), pseudotime analysis was performed to map developmental trajectories from day 0 to day 3 (**Figure 3C**). We next used CellOracle, which uses both scATAC-seq and scRNA-seq to predict gene regulatory networks that may be driving developmental transitions (Kamimoto et al., 2023). As internal validation of the method, in silico modeling predicted that genetic ablation of SP5, a known WNT negative regulator, would promote differentiation with a positive perturbation score along the developmental trajectory of Day 0, Day 1.5, and Day 3 cells (**Figure S3A**) (Huggins et al., 2017).

**Figure 3.**
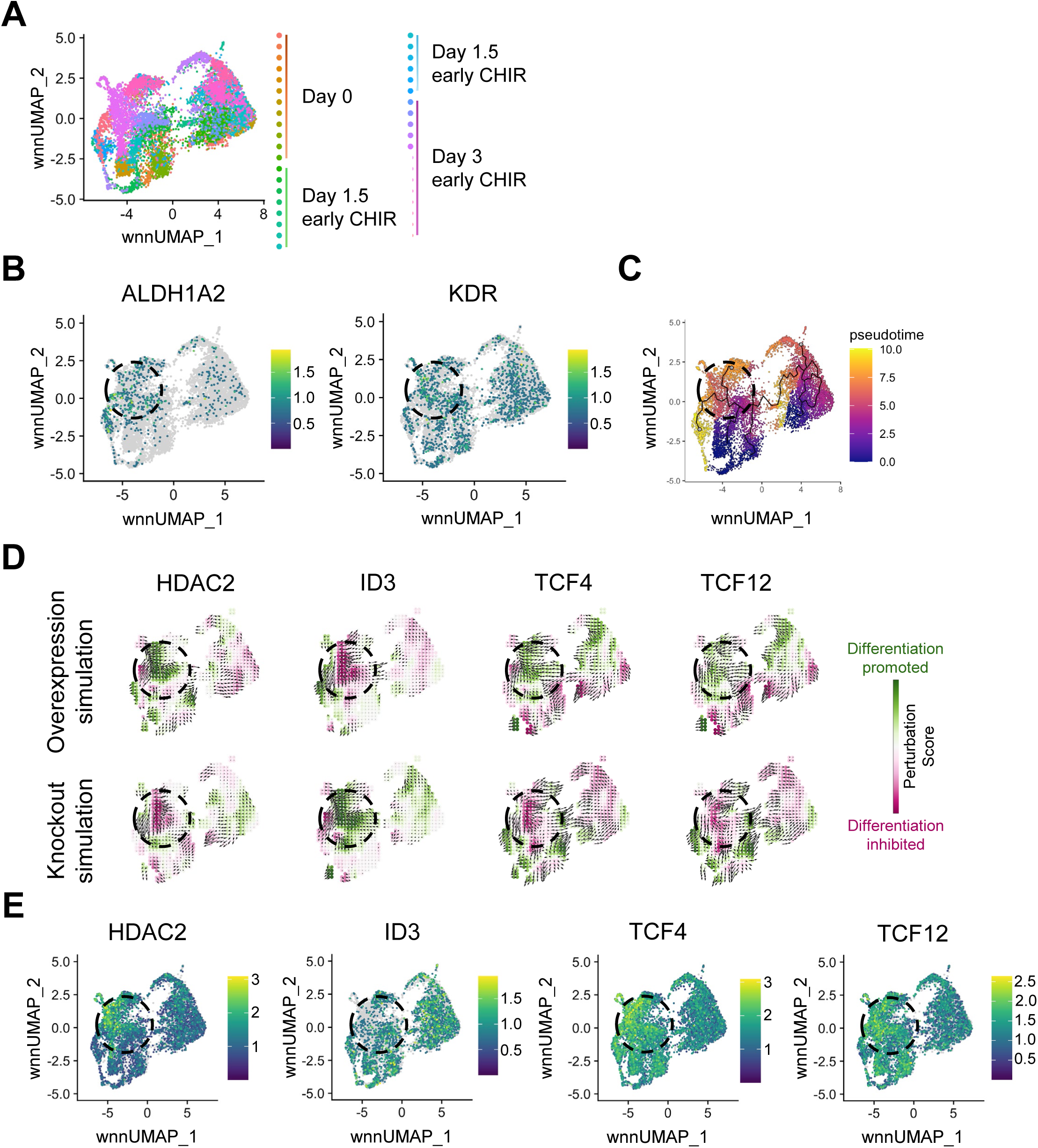
In silico gene perturbations identified transcriptional and epigenetic regulators in the development of *ALDH1A2*+ *KDR*+ RA-responsive mesodermal progenitors. (A) UMAP plot showing integrated scATAC- and scRNA-seq data from day 0 (undifferentiated), day 1.5, and day 3 of EB differentiation treated with CHIR99021 on day 0 (early CHIR). (B) Feature plots of ALDH1A2 and KDR relative expression to localize RA-dependent mesodermal progenitors on the UMAP embedding. Dashed circles represent cells enriched in ALDH1A2 and KDR. (C) Pseudo-time analysis using Monocle3 visualized on the same UMAP. Black lines indicate inferred differentiation trajectories from undifferentiated pluripotent stem cells toward differentiated RA-responsive mesodermal progenitor populations. (D) CellOracle in silico simulation to predict the effects of simulated overexpression and knockout of HDAC2, ID3, TCF4, and TCF12 on differentiation trajectories. Colors represent Perturbation Score, indicating predicted promotion (green, positive Perturbation Score) or inhibition (magenta, negative Perturbation Score) of differentiation upon simulated gene perturbation. (E) Relative expression of HDAC2, ID3, TCF4, and TCF12 in UMAP space.

We next examined other regulators also identified in our CellOracle analysis. ID3, TCF4, and TCF12 were predicted to strongly influence the transcriptional programs of Day 1.5 cells from Day 0 cells, while HDAC2 was predicted to be important for Day 3 cells to develop from Day 1.5 cells (**Figure S3B**). ID3, TCF4 (E2-2), and TCF12 (HEB) function as part of the ID–E protein network, in which ID proteins sequester TCF factors (E proteins) to regulate proliferation and lineage commitment, particularly during T and B cell development (Lazorchak et al., 2006; O’Riordan and Grosschedl, 1999; Roels et al., 2022). HDAC2 removes acetyl groups from histones, driving chromatin compaction and transcriptional repression, and has been shown to facilitate the endothelial-to-hematopoietic transition after mesodermal and hemogenic endothelial specification (Thambyrajah et al., 2018). In silico simulations predicted that TCF4, TCF12, and HDAC2 overexpression (OE) and ID3 knockout (KO) would promote differentiation toward Day 3 ALDH1A2+ KDR+ mesodermal progenitors (**Figure 3D**).

To consider the potential validity of these predictions, we examined the gene expression of these potential regulators throughout our scRNA-seq data. HDAC2 was highly expressed in the Day 3 ALDH1A2+ and KDR+ clusters, while ID3 was mostly absent in these cells. TCF4 and TCF12 expression remained largely unchanged during the differentiation process (**Figure 3E**). Together, these data predict a regulatory network in which TCF4, TCF12, ID3, and HDAC2 act as potential early drivers in the development of WNT-induced ALDH1A2+ KDR+ mesodermal progenitors.

### Functional validation confirmed HDAC2 as a key regulator required for RA-responsive mesodermal progenitor differentiation

To validate the functional roles of potential regulators in RA-responsive mesodermal differentiation identified through our CellOracle analysis, we performed CRISPR/Cas9-mediated knockouts of each gene and examined the impacts on differentiation in H1 hESCs. We electroporated H1 cells with Cas9 and guide RNAs (gRNAs) targeting HDAC2, ID3, TCF4, TCF12 individually, or TCF4 and TCF12 in combination. We then subjected these edited cells to EB differentiation in the presence of CHIR99021 to generate RA-responsive mesodermal progenitors. On day 3, differentiated EBs were dissociated into single cells and sorted by fluorescence-activated cell sorting (FACS) into KDR+ ALDEFLUOR+ (AF+) and KDR+ ALDEFLUOR- (AF-) mesodermal progenitor populations. Genomic DNA was extracted from sorted populations, followed by PCR amplification and Nanopore sequencing to assess frameshift mutation (FS) frequencies in each sorted populations, compared to day 0 undifferentiated cells.

Flow cytometry analysis revealed that knockout cultures of HDAC2 subtly but consistently showed decreased percentages of KDR+ AF+ mesodermal progenitors compared to the control (B2M knockouts) (**Figure 4A-B**), corroborating our prediction that HDAC2 promotes differentiation of these progenitors. In contrast, knockouts of ID3, TCF4, TCF12, or TCF4 and TCF12 did not lead to statistically significant differences in differentiation compared to control cells. As an additional way to quantify the impact of these KOs, we quantified frameshift mutation frequencies in sorted populations and compared with those in undifferentiated Day 0 cells (**Figure 4C**). For HDAC2 knockout cultures, Day 3 KDR+ AF+ cells had significantly lower frameshift mutation frequencies compared to both Day 3 KDR+ AF- and Day 0 starting cells, consistent with selection against cells that lack HDAC2. No such statistically significant changes were observed in knockout cultures of B2M, ID3, TCF4, TCF12, or TCF4 and TCF12.

**Figure 4.**
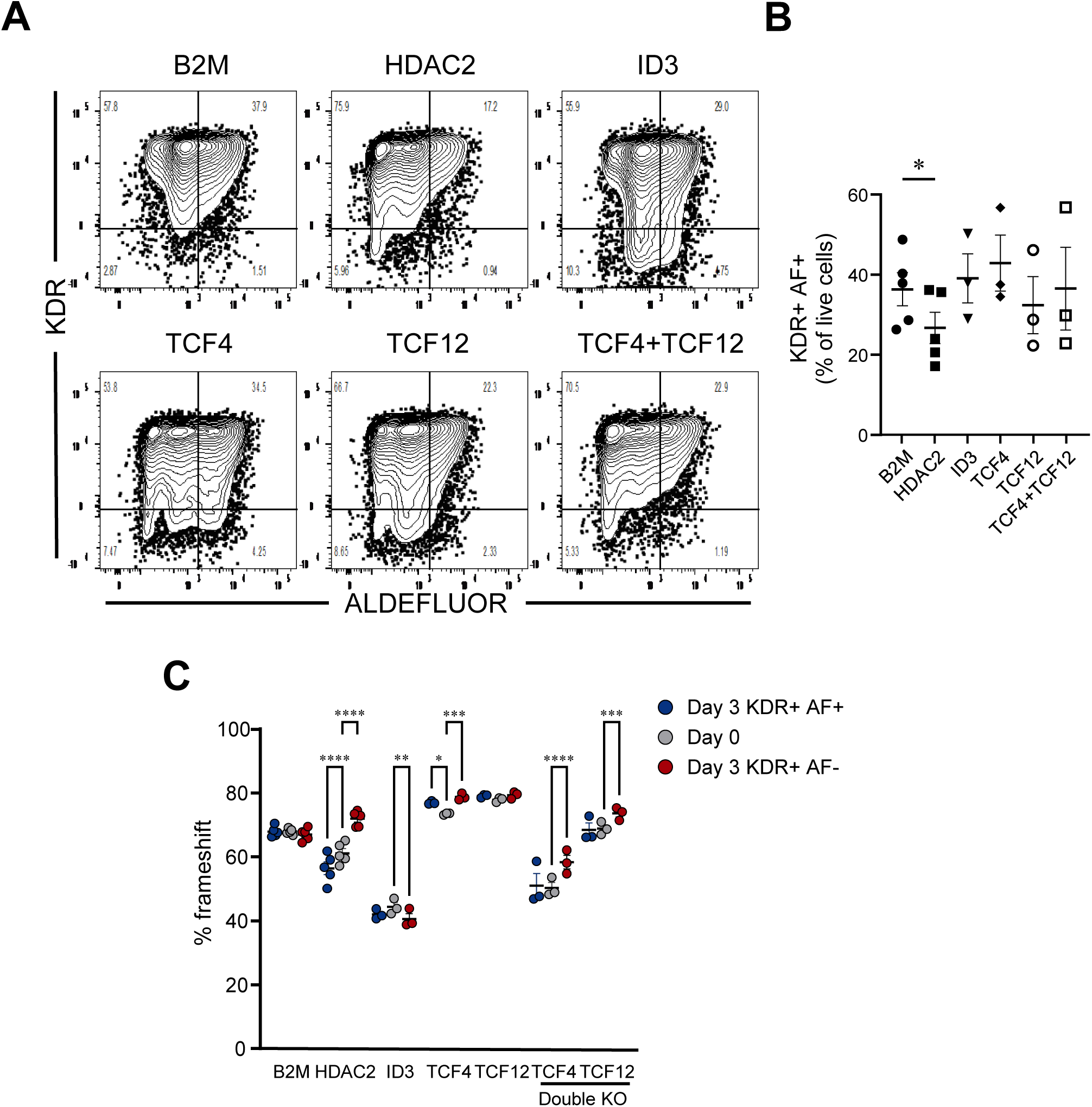
Functional validation confirmed HDAC2 as a key regulator required for RA-responsive mesodermal progenitor differentiation. (A) Representative flow cytometry plots showing the differentiation of KDR+ AF+ mesodermal progenitors on day 3 of differentiation. H1 hESCs were electroporated with Cas9-gRNA complexes targeting HDAC2, ID3, TCF4, TCF12 individually, or TCF4 and TCF12 in combination prior to EB differentiation. After 3 days, differentiated cells were sorted into KDR+ ALDEFLUOR+ (AF+) and KDR+ ALDEFLUOR- (AF-) populations via fluorescence-activated cell sorting (FACS). Genomic DNA was extracted, PCR-amplified, and sequenced using Nanopore sequencing to evaluate frameshift frequency. (B) Quantification of KDR+ AF+ mesodermal progenitor percentage. Statistical analyses were performed using 1-way ANOVA with Dunnet multiple comparisons test. *p < 0.05. Data are presented as mean ± SEM. Each symbol represents the value from an independent experiment. (C) Quantification of CRISPR/Cas9 gene-editing frameshift frequencies. Statistical analyses were performed using paired 2-way ANOVA with Šídák multiple comparisons test. *p < 0.05, **p < 0.01, ***p < 0.001, ****p < 0.0001. Data are presented as mean ± SEM. Each circle within a column represents the value from an independent differentiation.

For further validation, we overexpressed HDAC2 in H1 hESCs using mCherry-containing lentiviral vectors and examined how differentiation is subsequently affected. Compared to controls, HDAC2 overexpression promoted AF+ cell differentiation in the presence of CHIR99021 (**Figure 5A-B**). Additionally, we quantified the percentages of mCherry+ cells in AF+ populations and compared to those in AF- cells. In HDAC2 overexpression cultures, AF+ cells had subtly but statistically significantly higher mCherry+ percentages compared to AF- cells, indicating selection for cells that have HDAC2 overexpressed (**Figure 5C and 5D**). Together, these results confirmed that HDAC2 marks and promotes differentiation of ALDH1A2+ RA-responsive mesodermal progenitors. We did not observe statistically significant effects of ID1 and ID3 transcription factor overexpression on AF+ progenitors (**Figure S3A-B**).

**Figure 5.**
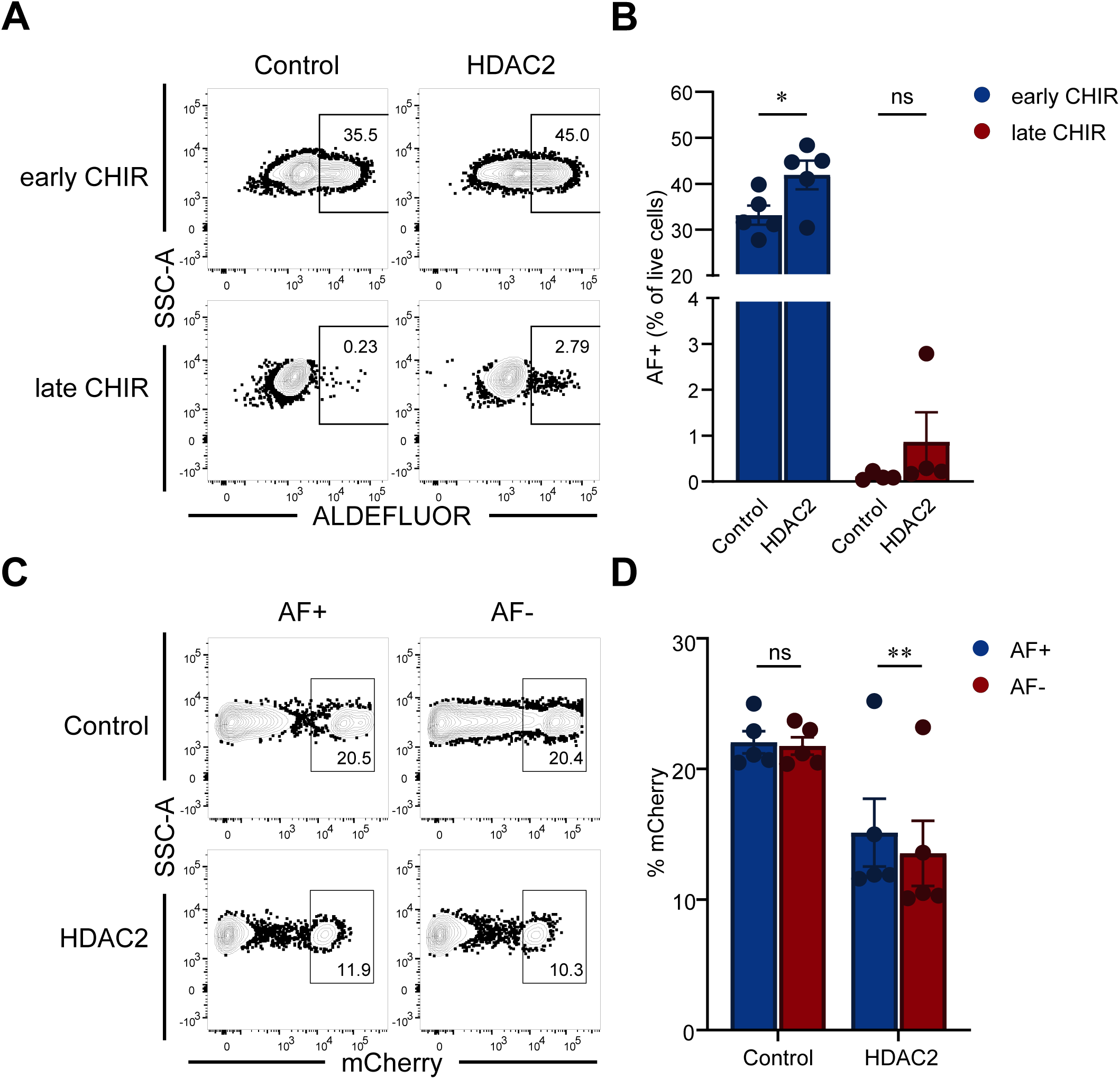
HDAC2 overexpression improves ALDEFLUOR+ progenitor differentiation. (A) Representative flow cytometry plots showing the differentiation of ALDEFLUOR+ (AF+) progenitors on day 3 of differentiation. H1 cells were transduced with mCherry-expressing lentiviral vectors encoding HDAC2, or control (empty vector). Transduced cells were then subjected to the same differentiation protocol as above with CHIR99021 on day 0 (early CHIR) or on day 2 (late CHIR). (B) Quantification of AF+ progenitor percentage. Statistical analyses were performed using paired 2-way ANOVA with Šídák multiple comparisons test. *p < 0.05. Each circle within a column represents an independent differentiation. (C) Representative flow cytometry plots quantifying mCherry+ cells within AF+ and AF- progenitors. (D) Quantification of mCherry+ percentage. Statistical analyses were performed using paired 2-way ANOVA with Šídák multiple comparisons test. **p < 0.01. Data are presented as mean ± SEM. Each circle within a column represents an independent differentiation.

### Drug screening revealed common histone modifications as epigenetic regulations in the development of RA-responsive mesodermal progenitors

As an orthogonal method to identify epigenetic factors that regulate the differentiation of ALDH1A2+ mesodermal progenitors, we conducted a chemical screen using a 152 compound library of small-molecule epigenetic inhibitors. EBs were treated with individual compounds at 10 μM final concentrations on day 0 and subjected to CHIR99021-induced mesodermal induction. On day 3 of differentiation, ALDEFLUOR+ cell numbers were quantified by flow cytometry and normalized to DMSO-treated controls. Most compounds had no significant impact on ALDEFLUOR+ cell differentiation (**Figure 6A**). However, CAY10683 and BG45, two known HDAC2 inhibitors, reduced the frequency of AF+ cells, corroborating the role of an HDAC2-dependent epigenetic program in promoting AF+ cells (**Figure 6A-B).** On the other hand, the top drug that enhanced the numbers of AF+ cells (**Figure 6C**) was GSK-LSD1, an inhibitor of histone demethylation, highlighting an additional epigenetic mechanism that supports AF+ cell formation. Together, these data demonstrate that early WNT activation during mesoderm induction promotes RA-dependent progenitors in part through epigenetic programs, such as those downstream of HDAC2 and LSD1.

**Figure 6.**
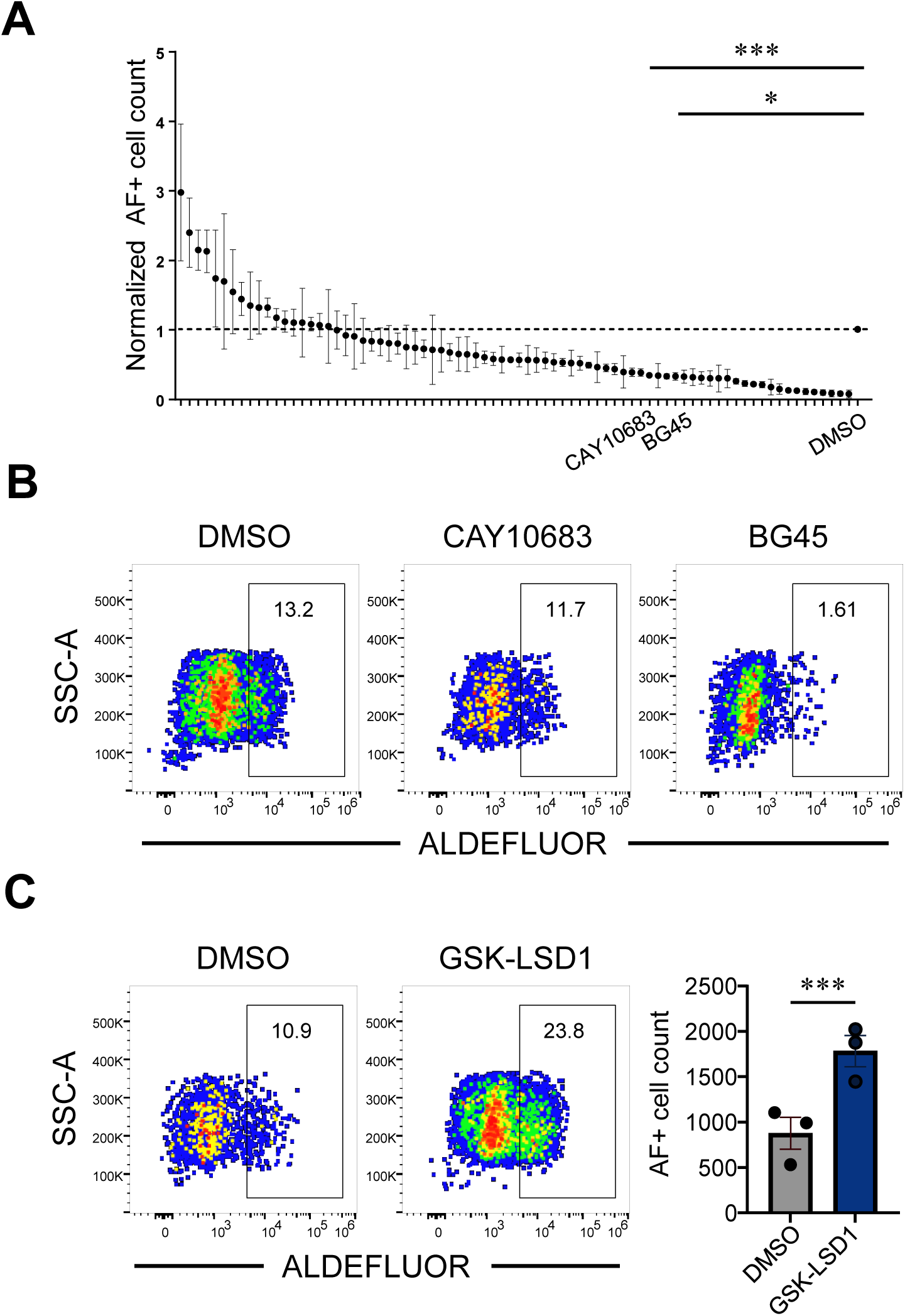
Drug screening revealed common histone modifications as epigenetic regulations in the development of RA-responsive mesodermal progenitors. (A) Quantification of ALDEFLUOR+ (AF+) cells in day 3 EB differentiation cultures treated on day 0 of differentiation with 10mM compounds from a library of 152 epigenetic small molecules. Values are normalized to the DMSO treated controls. Statistical analyses were done using 1-way ANOVA with Dunnet multiple comparisons test and *p < 0.05, ***p < 0.001. Data are presented as mean ± SEM from technical replicates. (B) Representative flow cytometry plots showing ALDEFLUOR expression in day 3 cultures treated with DMSO (control) and two HDAC2 inhibitors, CAY10683 and BG45. (C) Representative flow cytometry plots showing ALDEFLUOR expression after treatment with the top drug inhibitor GSK-LSD1 and DMSO (left panels). Quantification of absolute AF+ cell numbers across cultures treated with GSK-LSD1 and DMSO (right).

## Discussion

The specification of definitive hematopoietic progenitors from hPSCs has remained a barrier for adoptive cell therapies. Recent studies have shown that WNT pathway activation is critical to guide cells down the correct initial developmental pathways that lead to RA-responsive mesoderm and subsequent definitive hematopoiesis (Luff et al., 2022; Ng et al., 2024; Sturgeon et al., 2014). Here, we investigated pathways downstream of WNT signaling that mark and promote the formation of RA-responsive, ALDH1A2-expressing mesodermal progenitors. Gene regulatory network analysis predicted that both HDAC2- and E-box transcription factors were likely involved in RA-responsive mesoderm specification. Yet functional validation experiments were only able to confirm the role of HDAC2. It is possible that the E-box protein-dependent programs may not be necessary for the formation of these mesodermal progenitors, but may imprint subsequent lymphocyte potential. Studies in mice have shown that E-box transcription factors are critical for lymphopoiesis (Bain et al., 1994, 1997). Moreover, the reduction of E-box transcriptional programs may underlie age-dependent reductions in lymphopoiesis and function (Frasca et al., 2004a, 2004b).

We observed that HDAC2 promoted RA-responsive mesoderm, but only when WNT signaling was concurrently activated using CHIR99021. These data suggest that HDAC2 alone is insufficient to initiate RA-dependent mesodermal identity and instead functions in coordination with WNT-mediated transcriptional programs. One possible explanation is that HDAC2 facilitates the repression of lineage-inappropriate genes and perhaps also stabilizes chromatin accessibility at WNT target loci, thereby enhancing the transcriptional environment needed for ALDH1A2+ progenitor emergence. The impact of HDAC2 manipulation was relatively subtle, suggesting that partially redundant pathways may compensate for RA-responsive mesoderm specification. These findings highlight the importance of both epigenetic state and signaling context in regulating definitive mesodermal fate. For future studies, combining targeted chromatin modulation with stage-specific signals may improve differentiation efficiency.

Finally, our small molecule screen revealed that several distinct epigenetic modulators enhanced the generation of ALDH1A2+ mesodermal cells, including GSK- LSD1, which inhibits LSD1 demethylase activity. HDAC2 or HDAC1 can form complexes with LSD1 and coREST to regulate distinct transcriptional programs (Aufhauser et al., 2021). When HDAC2 is deleted, HDAC1-LSD1-coREST complexes become more stable, with HDAC1 directly promoting LSD1 demethylase activity (Aufhauser et al., 2021; Nalawansha and Pflum, 2017). This suggests a model in which HDAC2 promotes RA-responsive mesoderm, potentially by inhibiting LSD1 demethylase activity via a decrease in HDAC1-LSD1-coREST complex formation. Future studies should include integrating these chromatin-targeting agents into directed differentiation protocols, with the aim of selectively activating and silencing gene programs that favor the formation of ALDH1A2+ mesoderm. Studies such as ours show how modern single cell multiomic profiling can help guide these manipulations. These insights may accelerate the iterative optimization of protocols for definitive hematopoiesis and adoptive cell therapies.

## Experimental procedures

### Resource availability

#### Corresponding author

Further information and requests for resources and reagents should be directed to Deepta Bhattacharya.

### Materials availability

Plasmids and cells generated in this study are available upon request and completion of a materials transfer agreement.

### Data and code availability

Single cell multiome data is available at the NCBI Sequence Read Archive, Bioproject number PRJNA1272807.

### Generation of iPSC line

Tonsil samples were obtained from an individual undergoing elective tonsillectomy (Banner-University Medical Center). Tonsils were extracted from anesthetized individuals using Bovie electrocautery in a standard tonsillectomy surgery. Bilateral tonsils were subsequently combined and provided as a fresh sample for processing to the lab with no identifying information. 4 x 10^6^ CD19+ IgM+ were sorted and cultured in RPMI + 10% fetal calf serum for three days in the presence of 10 ng/ml of CD40L (Peprotech), 15 ng/ml IL- 2 (Peprotech), 10 ng/ml IL-6 (Peprotech), and 20 ng/ml IL-10 (Peprotech). Three hundred thousand CD19⁺IgM⁺ cells were infected with Sendai virus encoding OCT4, SOX2, KLF4, and MYC (CytoTune-iPS 2.0 Reprogramming Kit, Thermo Fisher Scientific) and cultured for three days in a xeno-free, blood-based medium on Matrigel (Churko et al., 2013). Cells were subsequently transferred to E7 medium for 12 days, followed by E8 medium until colony formation. Individual colonies were picked, clonally expanded, and cryopreserved at passage 3

### Cell culture and differentiation of hPSCs

H1 human embryonic stem cells (hESCs) and induced pluripotent stem cells (iPSCs) were maintained in feeder-free conditions on Matrigel-coated plates in mTeSR^TM^ Plus medium at 37°C with 5% CO2. Cells were passaged every three to four days using ACCUTASE. For differentiation, hPSCs were scraped off plates using Gentle Cell Dissociation Reagent and plated onto ultra-low attachment plates to form embryoid bodies (EBs). On day 0, differentiation medium made up of 73.5% IMDM, 25% Ham’s F12, 1% N2, 0.5% B27, and 0.05% BSA (Thermo Fisher) is supplemented with GlutaMAX (2 mM, Thermo Fisher), monothioglycerol (MTG; 0.45 mM, Sigma-Aldrich), ascorbic acid (50 ng/mL), transferrin (1.5 mg/mL, Sigma-Aldrich), bone morphogenetic protein 4 (BMP4; 10 ng/mL), basic fibroblast growth factor (bFGF; 5 ng/mL), and CHIR99021 (3 μM) as indicated. On day 2, SB-431542 (6 μM) was added. On the third day of differentiation, EBs were changed to StemPro-34 medium supplemented with GlutaMAX, MTG, ascorbic acid, transferrin, and bFGF, as above, with additional vascular endothelial growth factor (VEGF; 15 ng/mL) and all-trans retinal (RAL; 500 nM, Sigma-Aldrich). On day 6, interleukin 6 (IL-6; 10 ng/mL), insulin-like growth factor 1 (IGF-1; 25 ng/mL), interleukin 11 (IL-11; 5 ng/mL), stem cell factor (SCF; 100 ng/mL) and erythropoietin (EPO; 2 U/mL) were added. On day 10 and 13, Fms-related tyrosine kinase 3 ligand (FLT3L; 5 ng/mL) and interleukin 7 (IL-7; 5 ng/mL) were added. All differentiation cultures were maintained at 37 °C with 5% CO2. Unless indicated otherwise, all reagents and human recombinant factors were obtained from STEMCELL Technologies.

### Flow cytometry analysis

On day 3, EBs were dissociated into single cells using 0.25% Trypsin-EDTA for ten minutes. Cells were first stained for aldehyde dehydrogenase expression using the ALDEFLUOR™ Kit (StemCell Technologies, Cat# 01700) according to the manufacturer’s protocol. Control samples were established using diethylaminobenzaldehyde (DEAB), an ALDH inhibitor. Cells were then washed and stained in ALDEFLUOR™ Assay buffer with KDR–PE-Cy7 (clone 7D4-6, BioLegend, Cat# 359912, 1:100) for 45 minutes, and dead cells were excluded using DAPI staining. Cells were analyzed on an LSRFortessa (BD) cytometer. Fluorescence activated cell sorting (FACS) was performed on a 5-laser BD FACS ARIA II.

### T cell differentiation

Day 16 EBs were harvested and dissociated into single cells for fluorescence-activated cell sorting (FACS) using 0.25% Trypsin-EDTA for ten minutes, followed by Collagenase Type II for 30 minutes. A total of 500 to 2000 FACS-sorted CD34+ cells were added to individual wells of a 24-well plate coated with StemSpan™ Lymphoid Differentiation Coating Material (StemCell Technologies). Cells were cultured according to the manufacturer’s protocol for 14 days in StemSpan™ SFEM II medium supplemented withStemSpan™ Lymphoid Progenitor Expansion Supplement (StemCell Technologies). Following 14 days of differentiation, cells were analyzed using a BD-Fortessa flow cytometer (BD Biosciences). The antibodies used include CD13–PE-Dazzle™ 594 (clone WM15, BioLegend, Cat# 301720; 1:100), CD14–PE-Cy7 (clone M5E2, BioLegend, Cat# 301814; 1:100), CD45–FITC (clone 2D1, BioLegend, Cat# 368508; 1:100), CD5–PE-Cy5 (clone L17F12, BioLegend, Cat# 364031; 1:100), and CD7–PE (clone CD7-6B7, BioLegend, Cat# 343106; 1:100). Cells were sorted with a FACSAria II (BD) cell sorter and analyzed on an LSRFortessa (BD) cytometer.

### Combined single-cell ATAC-seq and RNA-seq

Single-cell suspensions were generated from Day 0 undifferentiated H1 hESCs, and Day 1.5 and Day 3 EBs, both with and without 3 μM CHIR99021 treatment. 10,000 nuclei per sample were processed and isolated using the Nuclei EZ Prep kit (Sigma-Aldrich) according to the manufacturer’s protocol. Libraries for both ATAC-seq and RNA-seq were prepared using the Chromium Next GEM Single Cell Multiome ATAC + Gene Expression Reagent Bundle (10x Genomics, PN-1000283). Libraries were prepared with i7 indices, multiplexed, and sequenced in partial lanes of the Illumina NovaSeq X Plus Series (PE150) to obtain 25,000 paired-end reads per nuclei. Unique sequences in each i7 index were then used for demultiplexing. Raw sequencing data from both ATAC-seq and RNA-seq were processed using Cell Ranger (10x Genomics) for alignment to the hg38 reference genome, filtering, and generating expression and accessibility matrices. These matrix files were then imported into RStudio for data processing which includes filtering out low quality cells, normalizing gene expression data using SCTransform and DNA accessibility data using latent sematic indexing (LSI). We then integrated all datasets into a single merged Seurat object following a previously described pipeline (https://stuartlab.org/signac/articles/pbmc_multiomic). We performed linear dimensional reduction on the integrated dataset to create Uniform Manifold Approximation and Projection (UMAP) graphs to visualize the clustering of cells based on both chromatin accessibility and gene expression profiles. Pseudotime analysis was then conducted using Monocle 3.0. Cells were ordered based on their differentiation status from undifferentiated pluripotent states (day 0) through intermediate (day 1.5) to differentiated progenitors (day 3). Cell trajectories were visualized on the UMAP embedding.

### Gene regulatory network (GRN) and perturbation modeling using CellOracle

CellOracle (v2.3.2) was used to infer transcription factor-driven regulatory networks. First, CellOracle identifies accessible regulatory elements (enhancers/promoters) from scATAC-seq data. Next, these regions are scanned for known transcription factor (TF) binding motifs, which define all possible TF-target gene interactions. CellOracle then integrates scRNA-seq data to retain only TF-target interactions supported by observed expression patterns. This results in dataset-specific gene regulatory networks (GRNs) that were used to identify TFs essential for development of a particular cell cluster. In silico perturbation simulations, overexpression (OE) and knockout (KO), were then used to predict regulatory impacts of a TF on differentiation trajectories. Perturbation Scores (PS) were calculated for each factor, quantifying their predicted ability to promote or inhibit differentiation.

### CRISPR/Cas9 gene editing of human embryonic stem cells

H1 hESCs were cultured under standard feeder-free conditions described above. 24 hours before electroporation, cells were moved to antibiotic-free mTeSR^TM^ Plus medium containing CloneR2 (Stem Cell Technologies). Cells were then harvested and electroporated using the Neon Transfection System (Thermo Fisher) with ribonucleoprotein (RNP) complexes consisting of recombinant Cas9 protein (40 μM, QB3 MacroLab) and synthetic single-guide RNAs (sgRNAs; 160 μM, IDT) targeting HDAC2, ID3, TCF4, TCF12 individually, or TCF4 and TCF12 simultaneously (double KO). B2M- targeting sgRNAs served as a negative control (**Table S1)**. Electroporated cells were plated onto Matrigel-coated plates containing antibiotic-free mTeSR Plus medium supplemented with CloneR2 for 24 hours, then maintained as described above without CloneR2. Edited cells were then subjected to EB differentiation in the presence of CHIR99021 to generate mesodermal progenitors.

### Genomic DNA extraction, PCR amplification, sequencing, and CRISPR editing efficiency analysis

Genomic DNA from FACS-isolated populations (Day 3 KDR+AF+ and KDR+AF- cells) and Day 0 undifferentiated cells was extracted using the Genomic DNA Blood/Cultured Cell Kit (IBI Scientific IB47207). Gene-targeted regions were PCR-amplified using primers specifically designed around sgRNA target sites, producing amplicons between 200 and 1,000 bp (**Table S1).** PCR products were purified using the Gel/PCR DNA Fragments Extraction Kit (IBI Scientific IB47082) and sent to Plasmidsaurus for amplicon sequencing using the Oxford Nanopore platform. Quantification of frameshift mutation frequency at each CRISPR target site was performed using CRISPResso2 (Clement et al., 2019).

### Small-molecule inhibitor screen

H1 hESCs were seeded into 96-well ultra-low attachment plates to induce embryoid body (EB) formation in Day 0 differentiation medium as described above. Small-molecule inhibitors (Cayman Chemical 11076, 152 total small molecules) targeting histone acetyltransferases, deacetylases, methyltransferases, and demethylases were added to the cultures at 10 μM final concentration using the Biomek FX liquid handling system. DMSO was used as a vehicle control. On Day 3 of differentiation, EBs were dissociated into single cells and stained for ALDEFLOUR, as described above. Flow cytometric analysis was performed using the 96-well autosampler of the Attune cytometer.

### Lentivirus construct design, production, and transduction

HDAC2, ID1, and ID3 genes were each cloned without stop codons into lentiviral vectors downstream of the E1Fα promoter and in-frame with a downstream T2A-mCherry cassette. Lenti-X 293T cells (Takara Bio USA) were cultured at 37°C with 5% CO_2_ in DMEM with 10% FBS, nonessential amino acids, Glutamax, sodium pyruvate, and penicillin/streptomycin. Cells were transfected at approximately 80% confluency in 10 cm^2^ tissue culture plates using 30 μL GeneJuice Transfection Reagent (Sigma-Aldrich) with 5 μg of lentiviral vector, 3.25 μg psPax2 (Addgene 12260), and 1.75 μg VSV.G (Addgene 12259). Medium was changed 6 to 8 hours post-transfection, and viral supernatant was harvested 48 and 72 hours later. 12 mL of viral supernatant was mixed with 3mL of 25% polyethylene glycol 8000 (Sigma-Aldrich) in PBS and incubated overnight at 4°C. This mixture was centrifuged at 3,000 x g for 20 minutes. Supernatant was discarded and the pellet was resuspended in 100μL PBS. Aliquoted lentivirus was stored at −80°C until time of use. 5 to 10 μL was used to transduce one well of H1 hESCs in a six-well plate.

### Statistical Analysis

Statistical significance was evaluated using a t test (alpha = 0.05) in Figures 1C-1D. In all other figures, one-way analysis of variance (ANOVA, alpha = 0.05) was performed to test for statistical significance. Data were presented as mean ± standard error of the mean (SEM). Statistical significance was defined as *p < 0.05, **p < 0.01, ***p < 0.001, and ****p < 0.0001.

### Figures

Figures were created with BioRender.com.

## Acknowledgements

This work was supported by the NIH 5R01EB035491 (D.B. and C.M.S.), R01HL172940 (C.M.S.), T32AG058503 (B.Q.T.) and P30CA023074 (for the Flow Cytometry Shared Resource). This work was also supported by the Gates Foundation (INV-1206188, INV- 071091 and INV-002414), the Gootter-Jensen Foundation, and the BIO5 Institute. We thank the Flow Cytometry Core and Functional Genomics Core at the University of Arizona, and Dr. Anthony Bosco for guidance on bioinformatics pipelines.

## Author contributions

Conceptualization: D.B. and B.Q.T.; Funding Acquisition: D.B., C.M.S., and B.Q.T.; Methodology: B.Q.T., S.L., J.M.C., C.M.S., Investigation: B.Q.T.; Formal Analysis: B.Q.T., Supervision: D.B. and C.M.S.; Writing (original draft): B.Q.T.; Writing-review and editing: D.B., S.L., C.M.S., J.M.C.

---

**Figure S1 related to Figure 1.**
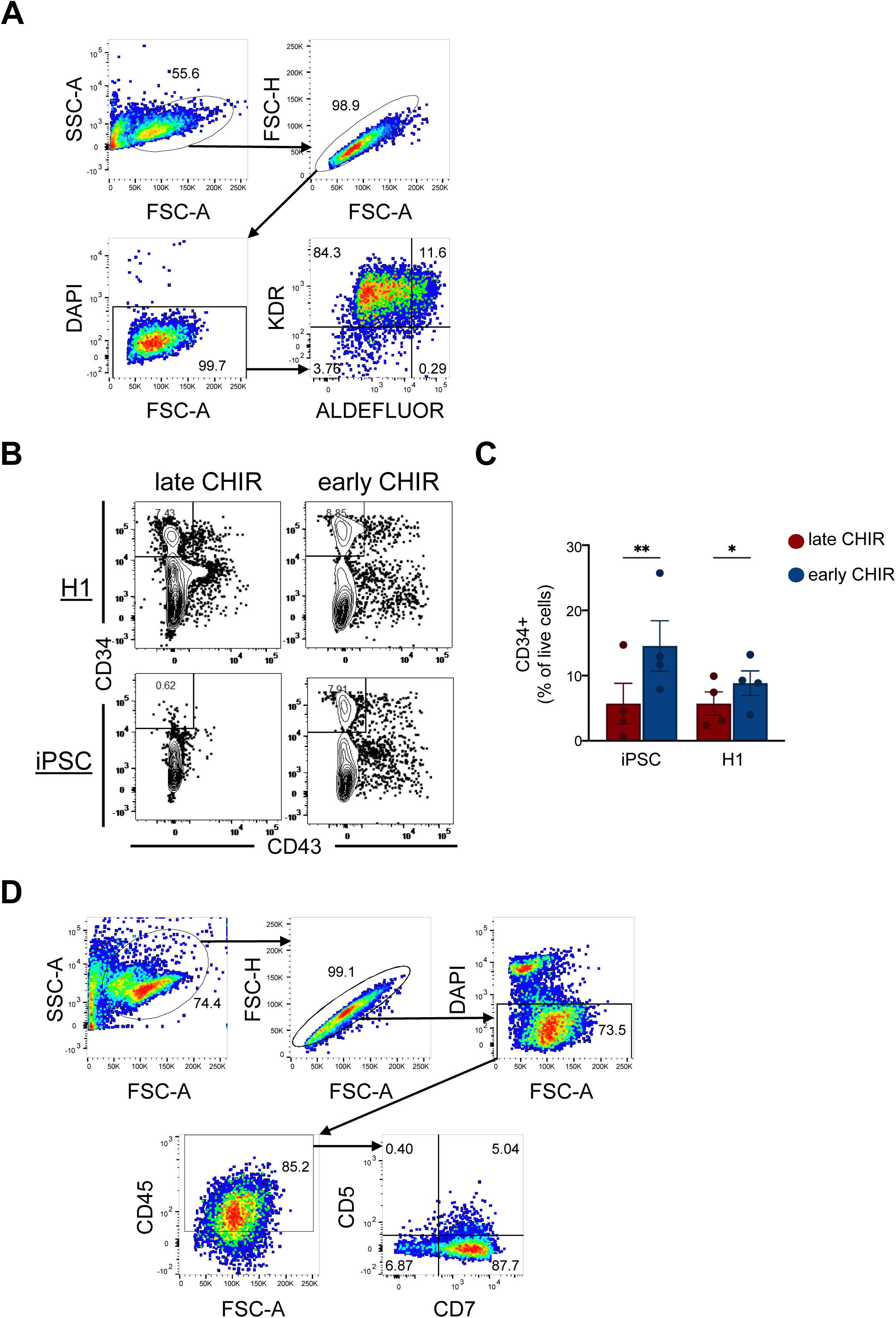
Differentiation of mesodermal progenitors from hPSCs and the role of CHIR 99021 in WNT signaling activation. (A) Gating strategy for flow cytometry analysis of KDR and ALDEFLUOR expression in H1- and iPSC-derived progenitors under early or late CHIR conditions. (B) Representative flow cytometry plots showing expression of CD34 and CD43 in H1- and iPSC- derived hematopoietic progenitors on day 16 of EB differentiation, with early or late CHIR treatments. (C) Quantification of the percentage of CD34+ cells in H1- and iPSC-derived cultures. Statistical analysis was performed using 2-way ANOVA with Šídák multiple comparisons test and *p < 0.05, **p < 0.01. Data are presented as mean ± SEM. (D) Gating strategy for flow cytometry analysis of CD5 and CD7 expression in H1- and iPSC-derived progenitors.

**Figure S2 related to Figure 2.**
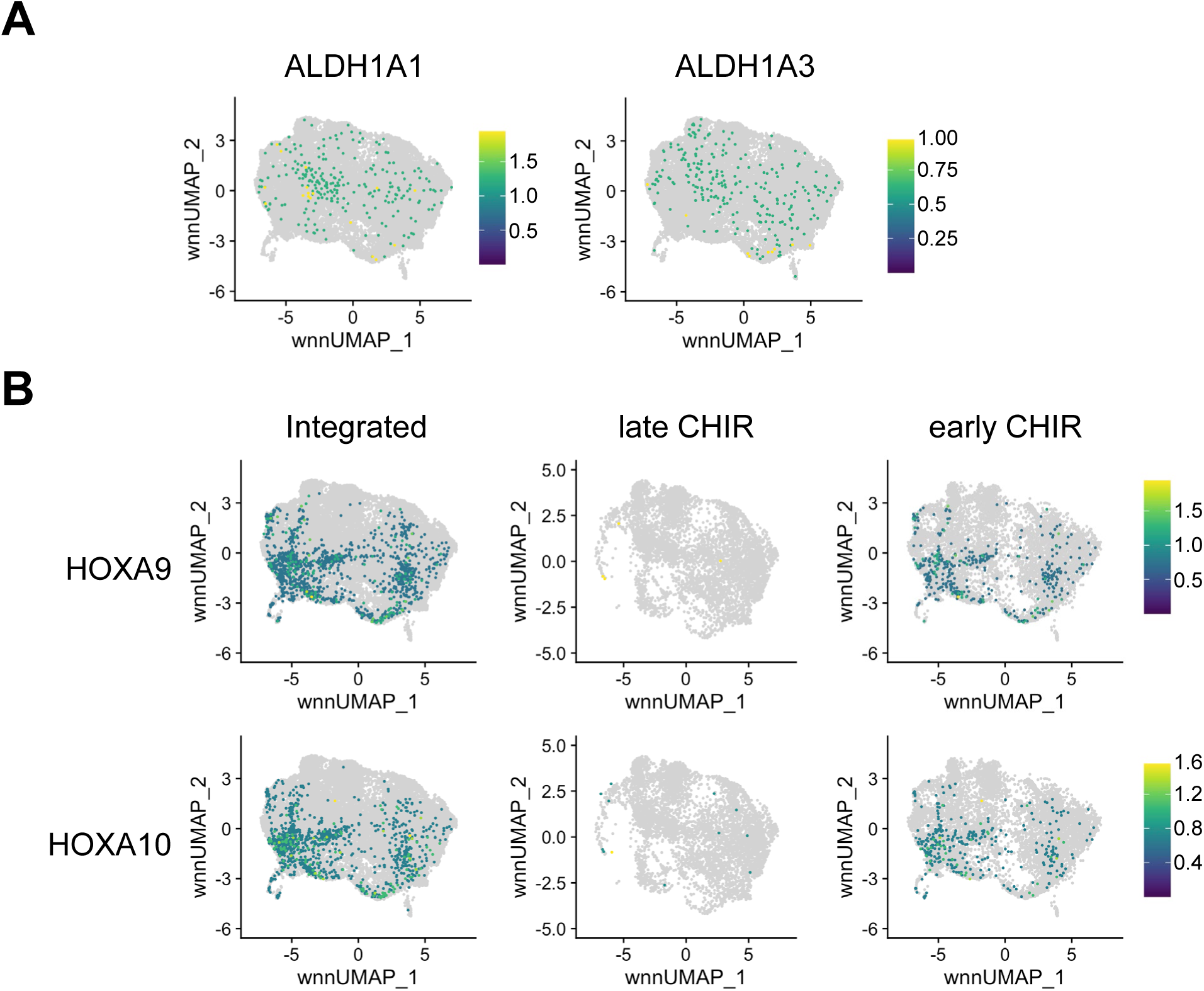
WNT signaling induces transcriptional changes associated with ALDH1A2- and HOXA-specific hematopoietic programs. (A) Relative expression of ALDH1A1 and ALDH1A3. (B) UMAP plots showing the relative expression of *HOXA9* and *HOXA10*.

**Figure S3 related to Figure 3.**
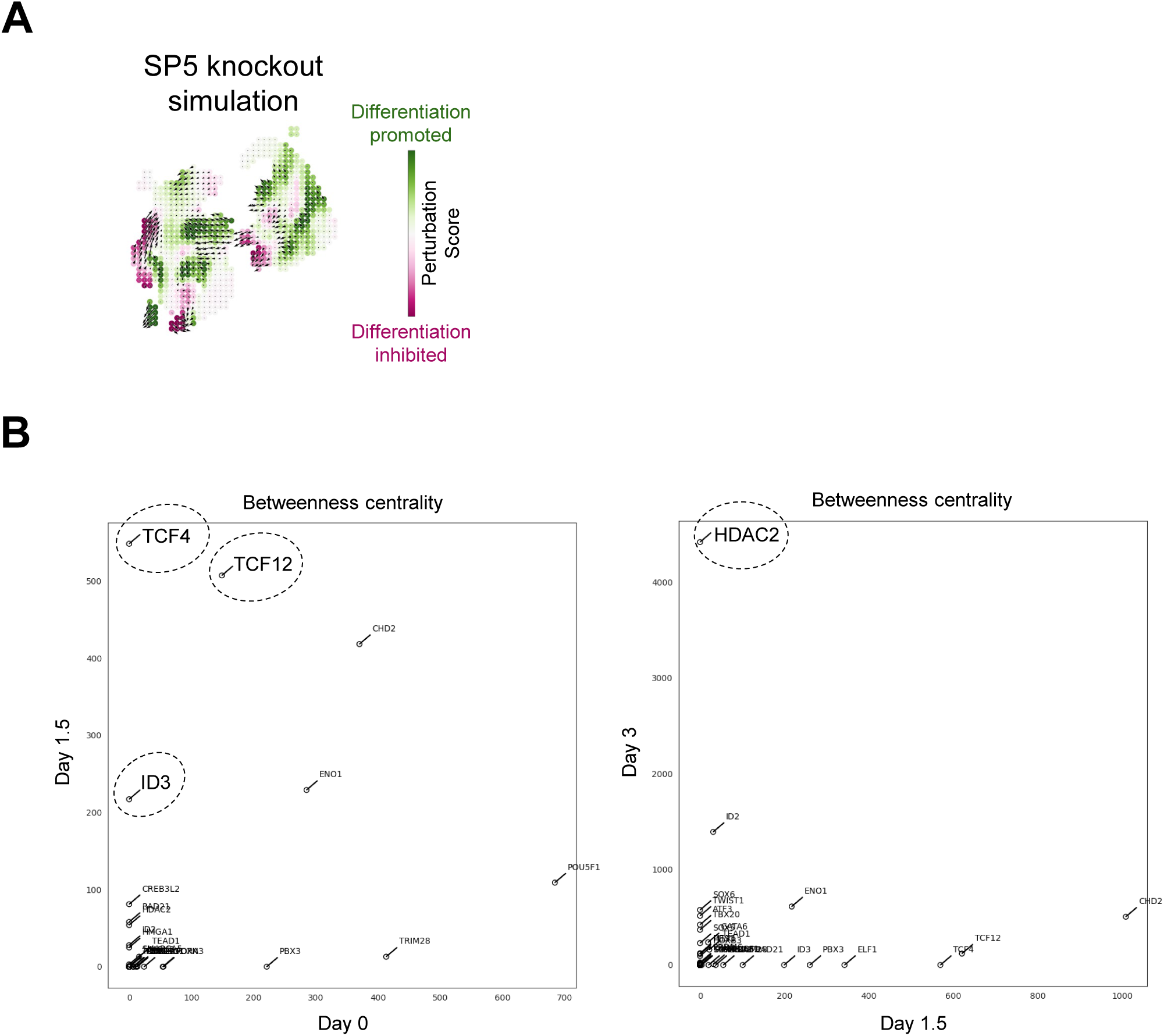
Identification of key regulators in RA-responsive mesodermal differentiation using CellOracle centrality analysis and in silico perturbation simulation. (A) CellOracle perturbation simulation showing the predicted effect of SP5 knockout on differentiation trajectories. Perturbation scores (PS) indicate where SP5 KO promotes (green, positive PS) or inhibits (magenta, negative PS) differentiation. (B) CellOracle network centrality analysis comparing TFs by betweenness centrality at Day 0 vs Day 1.5 cells (left) and Day 1.5 vs. Day 3 cells (right) of early CHIR cultures.

**Figure S4 relate to Figure 5.**
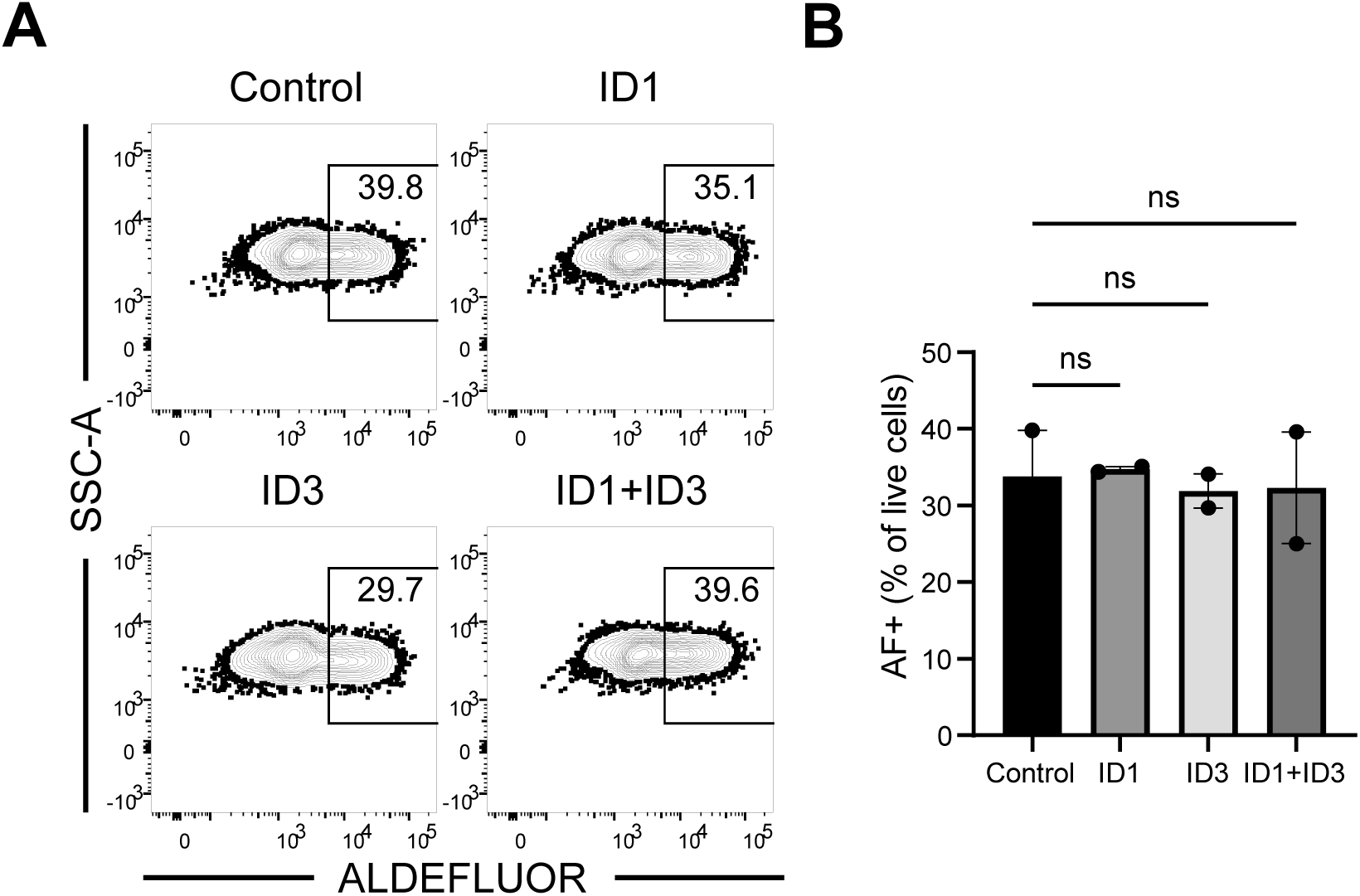
Overexpression of ID proteins does not impair generation of AF+ cells. (A) Representative flow cytometry plots showing the differentiation of ALDEFLUOR+ progenitors on day 3 of differentiation. H1 cells were transduced with mCherry-expressing lentiviral vectors encoding ID1, ID3, and ID1 and ID3 in combination, or control (empty vector). Transduced cells were then subjected to the same differentiation protocol as above with CHIR 99021. (B) Quantification of ALDEFLUOR+ progenitor percentage. Statistical analyses were performed using paired 1- way ANOVA with Dunnett multiple comparisons test. Data are presented as mean ± SEM.

**Table S1.**
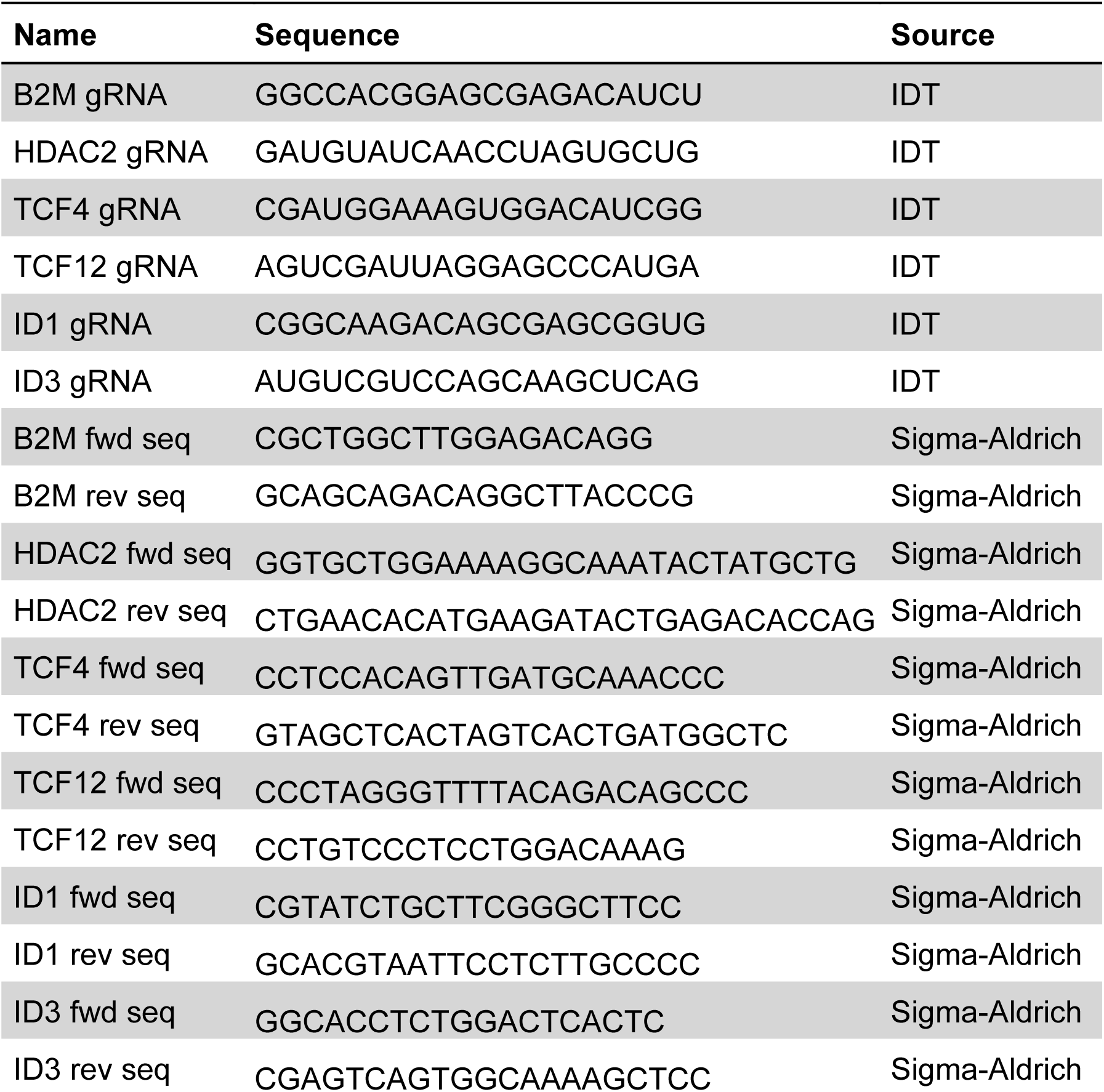
Sequences of guide RNAs and PCR primers to generate amplicons for Nanopore sequencing used in Cas9-knockout validation experiments.

